# Gender-specific brain activation during visual art perception

**DOI:** 10.1101/104166

**Authors:** Beatrice de Gelder, Rebecca Watson, Minye Zhana, Matteo Diano, Marco Tamietto, Maarten J. Vaessena

## Abstract

The human body is the most common object of pictorial representation in western art. The goal of this study was to probe its evolutionary basis of visual art perception by investigating neural markers of gender-specific brain activity triggered by paintings of male and female images. Our results show significant activity in brain areas other than those recently associated with visual arts perception. Novel findings concern participant-general as well as gender specific brain activity. Although our participants were fully aware that they were viewing artworks, the inferior parietal lobule - known for its role in the perception of emotional body images - and the somatosensory cortex – which is related to touch - were selectively active for female body paintings in all participants. The most interesting finding as regards gender was that the sight of female bodies activates the subgenual anterior cingulate cortex in males, an area known to subserve autonomic arousal. In contrast, in females the sight of the male body activated reward and control related parts of the dorsal anterior cingulate cortex. This supports the notion that basic evolutionary processes operate when we view body images, also when they are paintings far removed from daily experience.

## Introduction

Art has been around since the early dawn of mankind and the power in its images endures as a presence across all cultures (1, 2). Societies dedicate enormous financial resources and time to the creation and enjoyment of artworks. How do we understand the enduring attraction of art? Many have tried to answer this question, from traditional art historians to analysts of the currently booming art economy. Recently, neuroscientists have entered the discussion of human involvement in the arts by probing the perceptual basis of art, mainly in the visual arts and music. Interesting research findings cover a broad spectrum, ranging from visual analysis of artworks (3) to findings about motor resonance created in the viewer (4), and to inquiries into the neural basis of subjective aesthetic experience (5–7). Studies using functional magnetic resonance imaging (fMRI) appear to converge on medial orbitofrontal cortex, anterior cingulate and lingual gyrus as the major areas involved in visual art perception (5). Since our study uses images of paintings we expected those areas to play a role here. Furthermore, as we specifically selected paintings of bodies we predicted activity in areas in temporal cortex recently associated with perception of the body form and with movement and action perception mainly in parietal, premotor and somatosensory cortices (8, 9).

But aside from these, there may be so far undiscovered markers of the biological roots of visual art perception in the brain. This is suggested by Darwin’s explanation of the origins of art as well as broadly Similar to what has long been argued for the basic organization of sensory perception, cognition and behavior, art perception may have aspecific neural basis that is representative of its roots in the evolutionary history of the brain. Indeed, Darwin famously struggled with the manifestations of art and settled for a close link between art and its role in sexual selection. Now, adopting this general perspective challenges visual art perception studies to raise issues that are closely linked to processes of emotion, motivation and reward. This link has already been addressed by various authors, for example in the pleasure/reward/appetitive component of the aesthetic brain model (10, 11). Specifically, some studies using photographs have already advanced the evolutionary argument about art by looking at facial attractiveness from the perspective that physical beauty confers survival advantages (12, 13). This leads to the question whether similar preferences, possibly based on the evolutionary advantages conferred by attractiveness, can be found when whole body images are used. The specific hypothesis addressed here is whether beyond the previously reported brain areas involved in visual art perception there are neural markers of gender in brain activations as measured with functional MRI (fMRI) when people view classical paintings depicting male (“male paintings”) and female bodies (“female paintings”). Surprisingly, this hypothesis has not yet been tested with exemplars from the visual arts, specifically by using paintings that make up the bulk of the artistic environment western people are familiar with, in church decorations to cookie tins.

Our choice of materials was based on the following methodological requirements. First, we avoided imposing an aesthetic or judgmental experience frame of mind on the participants. Second, the images to be used needed to be broadly familiar in the sense that they were seen as traditional masterpieces of western painting in major museums, although any further specifics might be unknown to the participants. Paintings of the human body fit these constraints because they are universally seen as beautiful and recognized as artworks. To maximize homogeneity, paintings of the male body were selected from among representations of the San Sebastian theme. To render these images equivalent to the female set, the arrows were removed. Finally, to arrive at a balanced set for each gender, we created a group of female paintings with the arrows taken from the male paintings, see Fig. 1. In order to avoid any familiarity, memory or other cognitive processes related to the stimuli, a sample of naive college students was used.

**Figure 1.**
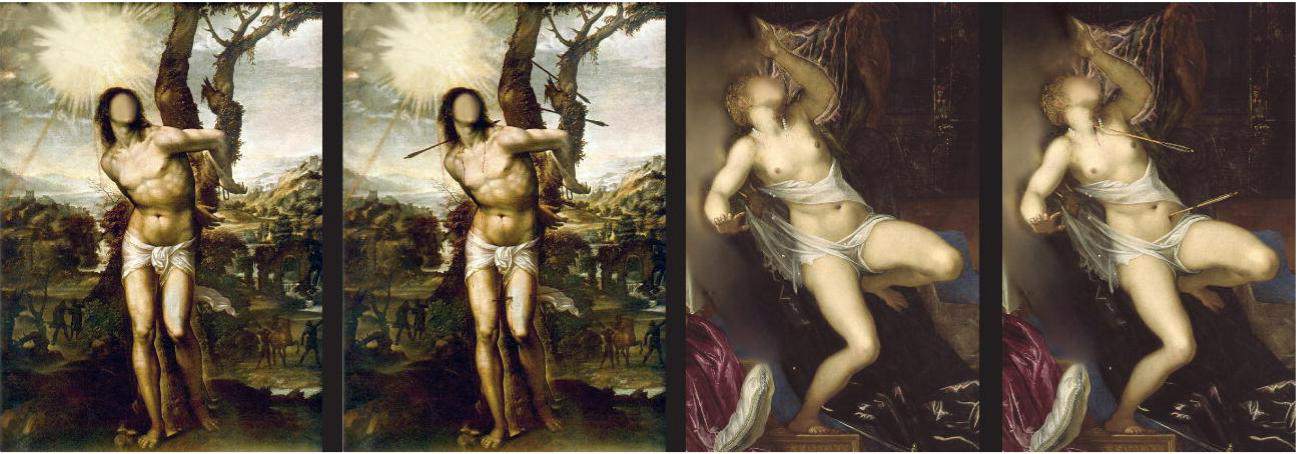
Example of stimuli. From left to right of male painting with no arrow, male with arrow, female with no arrow, female with arrow.

## Results

### Behavioral

Average looking times were calculated for each participant and each painting condition (Female with no arrow, female with arrow, male with no arrow, male with arrow) and submitted to a 2 x 2 (painting gender, arrow presence) repeated measures ANOVA. We observed a significant main effect of painting gender (F(1,15) = 6.194, p<0.03) and arrow presence (F(1,15) = 22.24, p<0.0001), but no significant interaction between the two factors. Inspection of the main effects showed that all participants generally spent longer looking at female paintings than male paintings, and longer at the paintings with arrows than those without. As we shall see, this pattern is not reflected in the brain activation data.

## fMRI results

The fMRI analyses were performed separately for all participants, and for two subgroups within our sample (male and female participants).

### All participants

The contrast of *female vs. male paintings* showed activity in the right inferior parietal lobule, bilateral lingual gyrus and left precentral gyrus. Only one cluster in the left superior temporal gyrus showed a stronger response to male vs. female paintings, see Fig. 2.

**Figure 2.**
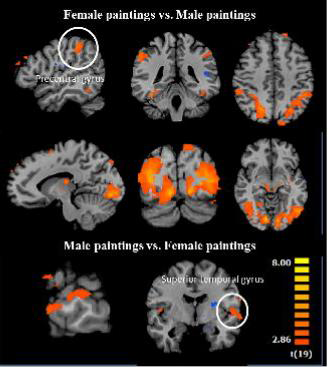
main results of the gender in paintings contrast. Top panel: increased activation for the female vs. male condition. Four significant clusters: peaks in the right inferior parietal lobule, the left precentral gyrus and the bilateral lingual gyrus. Bottom panel: increased activation for the male vs. female condition. One significant cluster: peak in the left superior temporal gyrus. Blue colors indicate negative t-values.

Next, within this contrast, the presence of *arrows vs no arrows* affects some activations differently. Paintings with no arrows, compared to those with arrows, elicited activity inthe bilateral medial frontal gyrus, left middle frontal gyrus, left precuneus and right parahippocampal gyrus. Paintings with arrows evoked activation in one cluster in the middle occipital gyrus, see Fig. 3.

**Figure 3.**
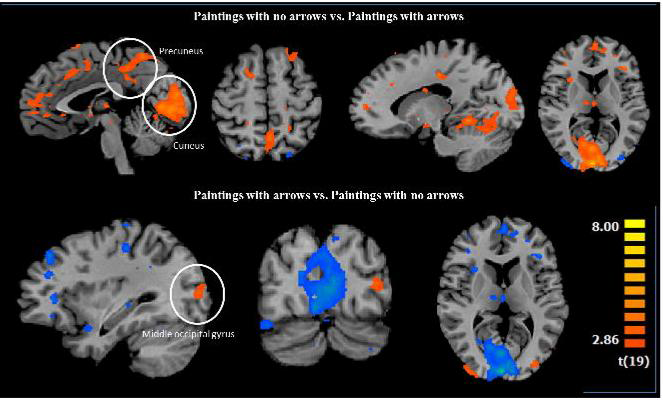
results for the arrows contrasts. Top panel: increased activations for the no arrow vs. arrow condition. Significant clusters in the right medial frontal gyrus and parahippocampal gyrus; cuneus; and left precuneus, medial frontal gyrus (two distinct clusters) and middle frontal gyrus. Bottom panel: increased activations for the reverse contrast. One significant cluster in the left middle occipital gyrus. Colors as in figure 2.

Finally, still for all participants we see that female paintings with no arrows vs. those with arrows showed activations in the bilateral superior frontal gyrus, bilateral medial frontal gyrus, right thalamus, left precuneus and left postcentral gyrus. No regions showed more activity to female paintings with arrows vs. no arrows. Finally, the contrast of male paintings with no arrows vs. male paintings with arrows evoked heightened responses in the right insula, cuneus and cingulate gyrus whereas the reverse contrast showed activity in the left middle occipital gyrus. Similarly, when we look at contrasts as a function of the gender of the paintings and the role of the arrows, we see that female paintings with no arrows vs male painting with no arrows revealed increased activation in middle frontal gyrus, fusiform gyrus, bi-lateral precentral gyrus, bilateral middle occipital gyrus, bilateral lingual gyrus and the right superior parietal lobule, see Fig. 4. For a complete overview of all participants results, see Table 1.

**Figure 4.**
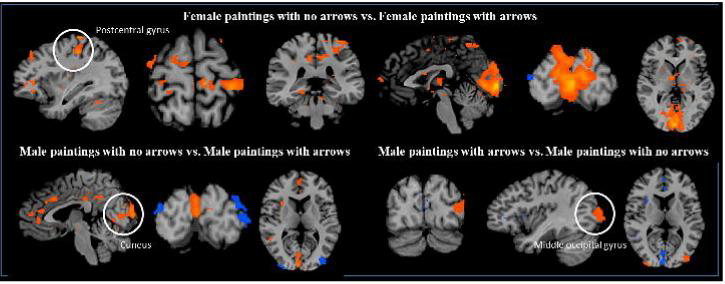
gender and arrow contrast results. Top panel: increased activations for the female-no arrow vs. female-arrow conditions. Significant clusters were found in the bilateral superior frontal gyrus, bilateral medial frontal gyrus, cuneus, right thalamus, and left precuneus (2 distinct clusters), postcentral gyrus and superior frontal gyrus. Bottom left panel: increased activations for the male-no arrow vs. male-arrow conditions. Four significant clusters in the claustrum, cuneus and cingulate gyrus (two distinct clusters). Bottom right panel: increased activations for the male-arrow vs. male-no arrow conditions. One significant cluster in the middle occipital gyrus. Color coding as in figures 2 and 3.

**Table 1:** results for the whole group GLM analysis.

| Brain Region (peak voxel) | Tailarach Coordinates |  |  | T Value | P value | Cluster size |
| --- | --- | --- | --- | --- | --- | --- |
|  | x | y | z |  |  |  |
| **Female paintings vs. Male paintings** |  |  |  |  |  |  |
| R Inferior parietal lobule | 51 | -37 | 43 | 4.49 | 0.000253 | 896 |
| R Lingual gyrus | 9 | -79 | -8 | 7.17 | 0.000001 | 31284 |
| L Lingual gyrus | -12 | -82 | -5 | 6.97 | 0.000001 | 34316 |
| L Precentral gyrus | -48 | -1 | 37 | 5.59 | 0.000022 | 965 |
| **Male paintings vs. Female paintings** |  |  |  |  |  |  |
| L Superior temporal gyrus | -63 | -7 | -2 | 4.73 | 0.000146 | 1686 |
| **Female paintings with no arrows vs. Male paintings with no arrows** |  |  |  |  |  |  |
| R Middle frontal gyrus | 57 | 32 | 17 | 4.95 | 0.000089 | 1185 |
| R Fusiform gyrus | 42 | -55 | -8 | 4.14 | 0.000554 | 1008 |
| R Precentral gyrus | 39 | 5 | 22 | 4.74 | 0.000143 | 1237 |
| R Middle occipital gyrus | 30 | -79 | 13 | 5.42 | 0.000032 | 6343 |
| R Superior parietal lobule | 33 | -52 | 55 | 4.64 | 0.000179 | 3886 |
| R Lingual gyrus | 9 | -79 | -2 | 6.40 | 0.000004 | 5545 |
| L Lingual gyrus | -12 | -82 | -5 | 7.51 | 0 | 14041 |
| L Inferior parietal lobule | -36 | -52 | 49 | 4.46 | 0.000271 | 894 |
| L Precentral gyrus | -51 | -1 | 34 | 5.46 | 0.000028 | 807 |
| **Male paintings with no arrows vs. Female paintings with no arrows** |  |  |  |  |  |  |
| **No significant voxels** |  |  |  |  |  |  |
| **Female paintings with arrows vs. Male paintings with arrows** |  |  |  |  |  |  |
| R Fusiform gyrus | 45 | -55 | -8 | 4.61 | 0.000192 | 771 |
| R Lingual gyrus | 9 | -82 | -5 | 6.63 | 0.000002 | 17555 |
| L Superior parietal lobule | -30 | -61 | 52 | 6.92 | 0.000001 | 23885 |
| L Inferior parietal lobule | -42 | -43 | 43 | 3.89 | 0.000987 | 729 |
| **Male paintings with arrows vs. Female paintings with arrows** |  |  |  |  |  |  |
| **No significant voxels** |  |  |  |  |  |  |
| **Paintings with no arrows vs. Paintings with arrows** |  |  |  |  |  |  |
| R Medial frontal gyrus | 18 | 26 | 31 | 4.57 | 0.000208 | 2228 |
| Cuneus | 0 | -91 | 10 | 7.11 | 0.000001 | 30975 |
| R Parahippocampal gyrus | 18 | -22 | -8 | 5.10 | 0.000064 | 792 |
| **L Precuneus** | -3 | -52 | 55 | 4.17 | 0.000519 | 2342 |
| **L Medial frontal gyrus** | -15 | 54 | 10 | 3.76 | 0.001332 | 1522 |
| **L Medial frontal gyrus** | -12 | 26 | 31 | 4.65 | 0.000173 | 1326 |
| **L Middle frontal gyrus** | -33 | 35 | 25 | 4.71 | 0.000153 | 806 |
| **Paintings with arrows vs. Paintings with no arrows** |  |  |  |  |  |  |
| **L Middle occipital gyrus** | -32 | -82 | 10 | 4.69 | 0.00016 | 696 |
| **Female paintings with no arrows vs. Female paintings with arrows** |  |  |  |  |  |  |
| **R Superior frontal gyrus** | 18 | 44 | 43 | 4.86 | 0.000108 | 2354 |
| **Cuneus** | 0 | -91 | 4 | 7.09 | 0.000001 | 30895 |
| **R Medial frontal gyrus** | 12 | 4 | 52 | 4.84 | 0.000113 | 1247 |
| **R Thalamus** | 6 | -4 | 4 | 5.45 | 0.00003 | 599 |
| **L Precuneus** | -6 | -49 | 52 | 5.23 | 0.000047 | 760 |
| **L Precuneus** | -9 | -34 | 43 | 5.13 | 0.00006 | 687 |
| **L Medial frontal gyrus** | -10 | 2 | 49 | 4.52 | 0.000235 | 937 |
| **L Superior frontal gyrus** | -9 | 65 | 28 | 4.51 | 0.000241 | 713 |
| **L Postcentral gyrus** | -33 | -31 | 64 | 5.38 | 0.000034 | 3725 |
| **L Superior frontal gyrus** | -24 | 26 | 55 | 5.70 | 0.000017 | 822 |
| **Female paintings with arrows vs. Female paintings with no arrows** |  |  |  |  |  |  |
| **No significant voxels** |  |  |  |  |  |  |
| **Male paintings with no arrows vs. Male paintings with arrows** |  |  |  |  |  |  |
| **R Claustrum** | 33 | 14 | 7 | 4.35 | 0.000348 | 745 |
| **R Cuneus** | 3 | -91 | 10 | 5.00 | 0.00008 | 1879 |
| **Cingulate gyrus** | 0 | -37 | 34 | 3.61 | 0.001882 | 772 |
| **R Cingulate gyrus** | 3 | 17 | 31 | 3.91 | 0.000939 | 728 |
| **Male paintings with arrows vs. Male paintings with no arrows** |  |  |  |  |  |  |
| **L Middle occipital gyrus** | -42 | -89 | 10 | 5.29 | 0.000042 | 2196 |

### Female participants

First we report the contrasts as a function of the gender depicted in the paintings. The contrast of female vs. male paintings showed activity in the right lingual gyrus, right middle occipital gyrus, right lingual gyrus and left middle temporal gyrus, left superior parietal lobule and left cuneus. No clusters showed a stronger response to male vs. female paintings. A detailed look inside this contrast shows that female vs. male paintings in the no arrow condition shows activity in right middle temporal gyrus and middle occipital gyrus. The same contrast in the arrows condition showed activation in right lingual gyrus, left cuneus and left cerebellum.

When we consider the overall contrast between no arrows vs. arrows, activation shows up in in the right precentral gyrus and cingulate gyrus, and left lingual gyrus. In more detail, the contrast of female paintings with no arrows vs. female paintings with arrows showed activations in the right fusiform gyrus and medial frontal gyrus, left cuneus and postcentral gyrus. No region showed more activity in female paintings with arrows. The contrast of male paintings with no arrows vs. male paintings with arrows evoked heightened responses in the right precentral gyrus, inferior frontal gyrus and bilateral anterior/mid cingulate gyrus, whereas the reverse contrast showed activity in the left middle occipital gyrus, see Fig. 4. For a complete overview of the female only participants results, see Table 2

**Table 2:** results for the female participant only.

| Brain Region (peak voxel) | Tailarach Coordinates |  |  | T Value | P value | Cluster size |
| --- | --- | --- | --- | --- | --- | --- |
|  | x | y | z |  |  |  |
| Female paintings vs. Male paintings |  |  |  |  |  |  |
| R Lingual gyrus | 36 | -67 | -2 | 5.38 | 0.000445 | 1316 |
| R Middle occipital gyrus | 30 | -79 | 19 | 6.73 | 0.000086 | 8849 |
| R Lingual gyrus | 9 | -79 | -2 | 7.23 | 0.000049 | 3244 |
| L Middle temporal gyrus | -39 | -79 | 22 | 7.61 | 0.000033 | 7028 |
| L Superior parietal lobule | -27 | -58 | 49 | 6.19 | 0.00016 | 2455 |
| L Cuneus | -24 | -82 | 28 | 4.76 | 0.001026 | 632 |
| Male paintings vs. Female paintings |  |  |  |  |  |  |
| No significant voxels |  |  |  |  |  |  |
| Female paintings with no arrows vs. Male paintings with no arrows |  |  |  |  |  |  |
| R Middle temporal gyrus | 42 | -67 | 16 | 12.68 | 0 | 23909 |
| L Middle occipital gyrus | -42 | -79 | 7 | 7.96 | 0.000023 | 15253 |
| Male paintings with no arrows vs. Female paintings with no arrows |  |  |  |  |  |  |
| No significant voxels |  |  |  |  |  |  |
| Female paintings with arrows vs. Male paintings with arrows |  |  |  |  |  |  |
| R Lingual gyrus | 9 | -79 | -2 | 6.44 | 0.00012 | 606 |
| L Cuneus | -21 | -79 | 19 | 5.59 | 0.000337 | 406 |
| L Cerebellum | -42 | -64 | -17 | 4.90 | 0.000848 | 427 |
| Male paintings with arrows vs. Female paintings with arrows |  |  |  |  |  |  |
| L Postcentral gyrus | -64 | -10 | 16 | 8.60 | 0.000012 | 1122 |
| Paintings with no arrows vs. Paintings with arrows |  |  |  |  |  |  |
| R Precentral gyrus | 45 | -1 | 7 | 7.44 | 0.00004 | 626 |
| L Lingual gyrus | -3 | -79 | 4 | 7.21 | 0.00005 | 7079 |
| R Cingulate gyrus | 9 | -1 | 37 | 5.37 | 0.000453 | 667 |
| Paintings with arrows vs. Paintings with no arrows |  |  |  |  |  |  |
| No significant voxels |  |  |  |  |  |  |
| Female paintings with no arrows vs. Female paintings with arrows |  |  |  |  |  |  |
| R Fusiform gyrus | 39 | -55 | -14 | 6.21 | 0.000157 | 807 |
| L Cuneus | -3 | -76 | 13 | 9.05 | 0.000008 | 13979 |
| R Medial frontal gyrus | 3 | -7 | 49 | 7.54 | 0.000035 | 2789 |
| L Postcentral gyrus | -36 | -22 | 46 | 6.55 | 0.000106 | 1005 |
| Female paintings with arrows vs. Female paintings with no arrows |  |  |  |  |  |  |

| No significant voxels |  |  |  |  |  |  |
| --- | --- | --- | --- | --- | --- | --- |
| Male paintings with no arrows vs. Male paintings with arrows |  |  |  |  |  |  |
| **R Precentral gyrus** | 45 | -1 | 7 | 7.37 | 0.000042 | 614 |
| **R Inferior frontal gyrus** | 33 | 5 | -14 | 5.62 | 0.000327 | 823 |
| **R Anterior cingulate** | 3 | 41 | 16 | 7.92 | 0.000024 | 2167 |
| **R Cingulate gyrus** | 3 | 17 | 34 | 5.53 | 0.000367 | 1063 |
| **L Cingulate gyrus** | -9 | -43 | 26 | 6.12 | 0.000175 | 848 |
| Male paintings with arrows vs. Male paintings with no arrows |  |  |  |  |  |  |
| **L Inferior occipital gyrus** | -30 | -79 | -2 | 5.38 | 0.000444 | 2002 |

### Male participants

The contrast of female vs. male paintings showed activity in the right middle frontal gyrus, right middle occipital gyrus, right precuneus, and left inferior occipital gyrus, left inferior parietal lobule and left inferior temporal gyrus. No clusters showed a stronger response to male vs. female paintings. In the sub-analysis we see that paintings with no arrows vs. arrows elicited activity in the cuneus, left anterior cingulate and thalamus.

When we consider the overall contrast between no arrows vs. arrows, the paintings with no arrows yield increased activation in cuneus, left anterior cingulate and left thalamus. Paintings with arrows (whether male or female) evoked activity in one cluster in the right middle occipital gyrus. The contrast of female paintings with no arrows vs. female paintings with arrows showed activations in the right lingual gyrus and left anterior cingulate and thalamus. The sub-contrast of female paintings with no arrow vs. arrows activated right lingual gyrus, left anterior cingulate and left thalamus. Left anterior cingulate was also seen for female paintings with arrows vs. no arrows, see Fig. 4. No activity was seen in the contrast of male paintings with no arrows vs. male paintings with arrows and vice versa. For a complete overview of the male participants results, see Table 3.

**Table 3:** results for the male participants only.

| Brain Region (peak voxel) | Tailarach Coordinates |  |  | T Value | P value | Cluster size |
| --- | --- | --- | --- | --- | --- | --- |
|  | x | y | z |  |  |  |
| Female paintings vs. Male paintings |  |  |  |  |  |  |
| R Middle frontal gyrus | 48 | 14 | 25 | 6.11 | 0.000177 | 639 |
| R Middle occipital gyrus | 30 | -79 | 10 | 5.70 | 0.000294 | 611 |
| R Precuneus | 18 | -61 | 43 | 4.95 | 0.000796 | 598 |
| L Inferior occipital gyrus | -33 | -85 | -2 | 9.02 | 0.000008 | 1905 |
| L Inferior parietal lobule | -33 | -55 | 37 | 5.24 | 0.000537 | 1185 |
| L Inferior temporal gyrus | -57 | -61 | -5 | 6.43 | 0.000121 | 1463 |
| Male paintings vs. Female paintings |  |  |  |  |  |  |
| No significant voxels |  |  |  |  |  |  |
| Female paintings with no arrows vs. Male paintings with no arrows |  |  |  |  |  |  |
| R Middle frontal gyrus | 51 | 14 | 25 | 6.21 | 0.000157 | 642 |
| Thalamus | 0 | -10 | 7 | 6.66 | 0.000092 | 658 |
| Male paintings with no arrows vs. Female paintings with no arrows |  |  |  |  |  |  |
| No significant voxels |  |  |  |  |  |  |
| Female paintings with arrows vs. Male paintings with arrows |  |  |  |  |  |  |
| R Cuneus | 21 | -79 | 28 | 6.42 | 0.000122 | 6497 |
| R Cerebellum | 21 | -70 | -11 | 6.79 | 0.00008 | 540 |
| L Inferior parietal lobule | -36 | -55 | 37 | 7.78 | 0.000028 | 4777 |
| L Middle temporal gyrus | -54 | -61 | -2 | 7.55 | 0.000035 | 6365 |
| Male paintings with arrows vs. Female paintings with arrows |  |  |  |  |  |  |
| No significant voxels |  |  |  |  |  |  |
| Paintings with no arrows vs. Paintings with arrows |  |  |  |  |  |  |
| Cuneus | 0 | -91 | 10 | 6.51 | 0.00011 | 2852 |
| L Anterior cingulate | -9 | 38 | -2 | 5.70 | 0.000294 | 646 |
| L Thalamus | -3 | -22 | 13 | 5.27 | 0.000513 | 696 |
| Paintings with arrows vs. Paintings with no arrows |  |  |  |  |  |  |
| R Middle occipital gyrus | 40 | -85 | 7 | 7.63 | 0.000032 | 575 |
| Female paintings with no arrows vs. Female paintings with arrows |  |  |  |  |  |  |
| R Lingual gyrus | 6 | -82 | -5 | 7.49 | 0.000037 | 3109 |
| L Anterior cingulate | -6 | 35 | -6 | 6.04 | 0.000194 | 808 |
| L Thalamus | 0 | -16 | 7 | 7.77 | 0.000028 | 2008 |
| Female paintings with arrows vs. Female paintings with no arrows |  |  |  |  |  |  |
| **L Anterior cingulate** | 33 | -85 | 4 | 5.13 | 0.000618 | 735 |
| **Male paintings with no arrows vs. Male paintings with arrows** |  |  |  |  |  |  |
| **No significant voxels** |  |  |  |  |  |  |
| **Male paintings with arrows vs. Male paintings with no arrows** |  |  |  |  |  |  |
| **No significant voxels** |  |  |  |  |  |  |

## Discussion

Our goal was to answer the question of whether there is evidence of sexual selection in the neural basis of classical western paintings of male and female bodies in participants with no knowledge or expertise in this domain. Our results show important and significant activity in brain areas beyond those recently associated with object based perception of the body, or with beauty and/or with reward, mainly in parietal, primary motor and somatosensory cortices. Furthermore, our results indicate a specific pattern of gender specificity suggesting that the preferential pattern seen here may reflect an evolutionary basis. We discuss this pattern now in more detail, first looking at the overall contrast for all participants between the presence or absence of arrows and then overall contrast between the gender of the images in the paintings.

Overall we see that all participants showed more activation for the no arrows than for the arrow condition suggesting that viewing pictures that may be related to pleasure constitutes a stronger trigger than those that may be related to pain. Importantly, besides posterior visual areas (cuneus and precuneus), the contrast no arrows-arrows activated the medial prefrontal gyrus (mPFC) bilaterally. The former are generally considered as visual areas although the precuneus has also been related to social processes (14) and is related to the fantasy content of visual materials (15). Activation of the mFPC is consistent with findings that the mPFC figures prominently in explanations of processes that have an affective component; furthermore, the medial orbitofrontal cortex and the ventromedial prefrontal cortex are activated when people judge objects to be beautiful (5, 16–19). Interestingly, the main activation in this contrast can be related to the pleasure/reward/appetitive component in the model by Chatterjee (10). The PFC is rich in sex hormone receptors, and has one of the highest concentration of oestrogen receptors in the human brain (20, 21). It is worth noting that contrary to what one might expect there was no increased activation triggered by the arrow condition in a number of areas related to pain perception (22, 23). This finding contrasts with the behavioral results showing that looking times are overall longer for images with than without arrows. This suggests that the presence of the arrows was noticed by the participants but did not trigger a brain reaction indicating pain related activity.

The female vs. male paintings contrast for all participants reveals inferior parietal, left precentral and bilateral lingual gyrus activity, areas that correspond to the sensorimotor component of the aesthetic experience (10). Inferior parietal lobule (IPL) activity was reported in a number of studies using body expressions (24–26) and is causally related to processing emotional body expressions (27). In this context it is interesting to note that these activations do not show in the contrasts by participant gender, except for IPL activation in the male participant group as discussed below.

To conclude the discussion of the two major contrasts, our results are consistent with the results of a meta-analysis of studies on visual aesthetics (6, 7, 28) in revealing a role for lingual gyrus, middle occipital gyrus, inferior and superior temporal gyri, precuneus and insula. Yet beyond that, there appears to be a major role for areas that have not come into the foreground in previous studies but that seem to have a clear significance here. These are mPFC, IPL and somatosensory cortex and their importance comes more to the foreground in the specific sub-contrasts.

We next consider the gender specificity in the no arrows condition for all participants. When looking more in detail at the female vs. male images in the no arrows vs. arrows contrast only, we see again a strong bilateral mPFC activity. This area is related to reward as well as to encoding beauty (16). Higher activity for female no arrow suggests that female bodies have higher reward value independently of the gender of the observer. Another aspect revealed in this sub-contrast is the activation of somatosensory cortex. Recent studies have shown that somatosensory cortex is reactive to not only external stimulation such as being touched as well as to mental imagery of touching (25), but also to the sight of body parts in situations of non-informative vision or visual enhancement of touch. Seeing a hand can enhance tactile acuity in the hand, even when tactile stimulation is not visible. Under normal conditions, touch observation activates the SI below the threshold for perceptual awareness (29). Vision of the body may act at an early stage in stimulus elaboration and perception, allowing an anticipatory tuning of the neural circuits in primary somatosensory cortex that underlie tactile acuity (30). Note, this effect is obtained for all participants underscoring that it is specific for this type of stimulus. A speculative interpretation may be that female bodies convey to the brain of the observer a tactile experience, in line with studies on touch showing that thalamus goes with primary somatosensory cortex (25, 31, 32).

Last, we discuss the relation between the gender of the participants and the paintings. The most intriguing aspect of our results is in the combined effects of gender of the participant and gender of the stimuli. Interestingly, gender-specific effects of stimulus type are found in one specific area, the anterior cingulate cortex (ACC). Here the effects are more specifically located in two different subsections of the ACC and appear to follow the dorsal-ventral division and functions of the ACC. Female participants show greater activation to male images in dorsal ACC (dACC) and male viewers more to female paintings in subgenual ACC (sgACC).

First, brain activations of female participants looking at male paintings revealed increased activation in the dACC. The dACC is thought to play a crucial role in the development of human cognitive control and guiding behavior (33). It is associated with attention modulation, competition monitoring, complex motor control, motivation, novelty, error detection, and the modulation of reward-based decision making (for review, see (34)). Meta-analyses of the neuroimaging literature have confirmed that the dACC plays a central role in control-demanding tasks (35–38). The role of dACC may be that of monitoring (39–41). Second, when male participants look at female paintings the pattern of activations reveals a different, equally specific activation, this time in sgACC. Interestingly, this area is often reported in relation to emotion and arousal. It has long been recognized that the sgACC contributes to autonomic control and the sgACC is densely interconnected with structures that play a central role in visceromotor control, such as the hypothalamus (for a review see (42)). The sgACC may contribute to positive affect by sustaining arousal in anticipation of positive emotional events (43). The sgACC is densely connected with mesolimbic pathways that facilitate the release of oxytocin (44) – a neuropeptide which bolsters interpersonal trust and cooperation (45) – and also sends direct projections to subcortical areas that control autonomic responses (46). Lesions in this area result in blunted responses to emotionally meaningful stimuli (47). Other studies have contrasted the different roles of sgACC and the dACC based on anatomical connectivity: a pre-genual region strongly connected to medial prefrontal and anterior midcingulate cortex and a subgenual region with strongest connections to nucleus accumbens, amygdala, hypothalamus, and orbitofrontal cortex (Johansen-Berg 2008). Interestingly, the connections between mPFC and amygdala have recently been viewed as targets for understanding the role of internalizing and psychopathology (48).

Previous studies have used paintings as a means to probe the neural basis of beauty perception and later findings have highlighted a multicomponent system consisting of emotion-valuation, sensorimotor and knowledge–meaning (10). Our findings about specific gender related effects are consistent with that literature but also extend and modify it significantly. Concerning the emotion-valuation and the appetitive component, we find that the mPFC activation is specific for female stimuli independent of the gender of the participants. The sensorimotor component, IPL and motor activation are each equally stimulus gender-specific.

On the other hand, our results represent an important step forward in understanding gender effects in artistic experience. Previous studies looking at gender effects in body perception have mainly focused on brain activation to neutral bodies in EBA and reported right lateralization in women (49) including when an aesthetic/liking of natural body appearance was measured (50). When emotional whole body expressions were studied gender/stimulus specific effects were found for male participants viewing male anger expressions (51).

In conclusion, our results show important and significant activity in brain areas beyond those recently associated with perception of the body, of subjective liking and of the experience of beauty in visual arts in general. What is common to both genders when viewing the female body are the IPL and somatosensory cortex. While our participants were fully aware that they were viewing artworks, the brain mobilizes IPL, known for its role in perception of emotional body images (8) and for its relation to embodiment (52). Similarly, somatosensory cortex activation associated with the sight of the female body experience are active, consistent with findings on emotions and tactile experience. On the other hand, the most specific finding concerning the role of gender is that the sight of female bodies activates in males an area known to subserve autonomic arousal. However, in females the sight of the male body activates reward and control related part of the dACC. Taken together, the general and the gender specific activities provide support for the notion that basic bodily experience processes operate when we view body images, regardless of the fact that they are artifacts.

## Materials and Methods

### Participants

Twenty healthy participants (10 males, mean age of the whole group 25y, range 21-29y) participated in the study. All had normal or corrected-to-normal vision, and no history of neuropsychiatric disorders. None of the participants had a previous background in art or had any special interest in painting. The experiment was approved by the Ethics committee of Maastricht University, and written informed consent was obtained from each participant beforehand. Participants were screened for fMRI experimentation safety and received monetary compensation.

### Stimuli

16 male and 16 female full-body classical oil painting were selected from the internet (for a full list of stimuli, see Supplementary Table 1). The theme of the male-body paintings was Saint Sebastian pierced with arrows, in either a standing or half-lying position. Female-body paintings were selected from the themes of Andromeda, Cleopatra, Danae and Venus, in standing or lying positions. Some male paintings showed the bodies bound with ropes or chains and these were modified in Photoshop CS6 (Adobe systems incorporated, USA) in order to create a new more homogeneous set for four different stimulus conditions: “male bodies with arrows”; “male bodies with no arrows”, “female bodies with arrows”; and “female bodies with no arrows”. Firstly, as the original male body paintings all depicted bodies pierced with arrows, the arrows and blood from the paintings (including those in the background) were removed to create the “male bodies with no arrows” set. Furthermore, as none of the original female body paintings showed bodies pierced with arrows, we created the “female with arrow stimuli” set by copying the arrows from the San Sebastian paintings, and adding 2-4 arrows with roughly matching painting style onto the limbs, the torso, or the neck of the female bodies, adding shadows and some blood drops accordingly. The number of arrows in the female paintings roughly matched the number in male paintings (Male: arrow on the bodies: mean=2.64, SD=1.28; arrow in the scene: mean=3.29, SD=2.20. Female: mean=2.75, SD=0.68). Finally, for each of the four stimulus sets the images were then cropped to contain only the body of interest. The faces were blurred for each body and all the other people or any background were also blurred. See Figure 1 for illustrative examples of the modified sets.

### Design

#### Behavioral

In an offline behavioral experiment, 13 new participants (10 female, mean age ± standard deviation 23 ± 6.1 years) were presented with each painting in turn, and instructed to inspect it for as long as they wanted to. Paintings were presented on a PC screen, using Presentation software (Neurobehavioural Systems, San Francisco, CA). When participants were ready to view the next painting, they pressed a key on the computer keyboard. Here the aim was to gain an offline measure of the relative looking time per image.

#### fMRI

Participants were scanned using a Siemens 3T Prisma scanner (Siemens, Erlangen, Germany). Earplugs were used to attenuate scanner noise and padding was used to reduce head movements. All stimuli presented during the fMRI session were projected to a clear screen at the back of the scanner bore that participants could see using a mirror mounted on top of a head coil. The study consisted of 4 experimental conditions: female paintings with arrows; female paintings with no arrows; male paintings with arrows; male paintings with no arrows. The experimental design was blocked, with ten blocks per condition (40 blocks in total) and 8 trials per block. Participants passively viewed the stimuli within each block, and were not instructed to perform any task. Each stimulus was presented in the center of the screen, on a white background. The shorter side of the stimuli was resized to 408 pixels, and the long side varied between 547 to 1084 pixels. All stimuli spanned within the visual angle of 19.02x19.49 degrees. The order of blocks was pseudo randomized; additionally the order of the trials within each block was randomized. Within blocks, each painting was presented for 1800ms, and the inter-trial interval was 200 msec. Time between blocks was 12000 msec. Stimuli were presented using Presentation software (Neurobehavioural Systems, San Francisco, CA).

#### MRI parameters and Functional Data Processing

Both high-resolution anatomical [T1-weighted, flip angle (FA) = 9 degrees, TR = 2250, TE = 2.6 msec, 192 slices, field of view (FoV) = 256 mm, isotropic voxel resolution of 1 x 1 x 1 mm3] and whole-brain functional images [T2*-weighted echo-planar imaging: TR = 2000, TE = 30 msec, 35 contiguous slices, slice thickness = 3 mm, voxel resolution = 3 x 3 x 3 mm3] were obtained. FMRI data were processed using BrainVoyager QX (Brain Innovation, Maastricht, The Netherlands). Pre-processing included slice acquisition time correction, temporal high-pass filtering, rigid-body transformation of data to the first acquired image to correct for motion, and spatial smoothing with a 4mm FWHM gaussian kernel. Functional data were co-registered to anatomical data per subject, and further transformed to Talairach space.

#### Activation Data Analysis

BOLD time courses of 16 sec individual blocks were regressed onto a pre-specified model in a conventional GLM. Separate predictors were implemented for the four different conditions. We then computed a group statistical map, calculated by using a random-effects (RFX) model, restricting this by using a mask to exclude non-brain matter voxels. Further to this, we computed the following t-contrasts: Female paintings vs. Male paintings; Paintings with no arrows vs. Paintings with arrows; Female paintings with no arrows vs. Female paintings with arrows; Male paintings with no arrows vs. Male paintings with arrows. We also computed these contrasts at the group level for female and male participants separately.

The statistical thresholding and multiple-comparison correction was performed in a two-step procedure. First, a single voxel threshold of p = 0.01 (uncorrected) was used for initial statistical maps. Next, a whole-brain correction criterion was calculated by estimating a false-positive rate for each cluster. This was established by means of Monte-Carlo simulation (1000 iterations), with the minimum cluster size threshold applied to the statistical maps corresponding to a cluster-level false positive rate (α) of 5%. Cluster size is reported in number of anatomical voxels.

**Figure 5.**
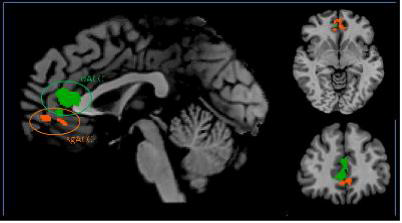
gender of participant effects. The ACC clusters found for the female paintings with no vs. with arrows, for male participants in the sgACC (orange) and male paintings with no vs. with arrows for female participants in the dACC (green).

## Acknowledgements

This work was supported by a European Research Council (ERC) grant under the European Union Seventh Framework Programme for Research 2007–2013 (grant agreement number 295673) and the European Union’s Horizon 2020 Research and Innovation Programme under grant agreement No 645553, ICT DANCE (IA, 2015-2017).

## References

1. Mithen SJ (1996) The Prehistory of the Mind a Search for the Origins of Art, Religion and Science.

2. Freedberg D (1991) The Power of Images: Studies in the History and Theory of Response (University of Chicago Press).

3. Cavanagh P (2005) The artist as neuroscientist. Nature 434(7031):301– 307.

4. Freedberg D & Gallese V (2007) Motion, emotion and empathy in esthetic experience. Trends in cognitive sciences 11(5):197– 203.

5. Ishizu T & Zeki S (2011) Toward a brain-based theory of beauty. PLoS One 6(7):e21852.

6. Ishizu T & Zeki S (2014) A neurobiological enquiry into the origins of our experience of the sublime and beautiful. Frontiers in human neuroscience 8:891.

7. Zeki S (2011) Splendors and miseries of the brain: Love, creativity, and the quest for human happiness (John Wiley & Sons).

8. de Gelder B (2016) Emotions and the Body (Oxford University Press, Incorporated).

9. De Gelder B (2006) Towards the neurobiology of emotional body language. Nature Reviews Neuroscience 7(3):242– 249.

10. Chatterjee A & Vartanian O (2014) Neuroaesthetics. Trends in cognitive sciences 18(7):370– 375.

11. Chatterjee A (2013) The Aesthetic Brain: How We Evolved to Desire Beauty and Enjoy Art (OUP USA).

12. Hahn AC & Perrett DI (2014) Neural and behavioral responses to attractiveness in adult and infant faces. Neuroscience & Biobehavioral Reviews 46:591– 603.

13. Aharon I , et al. (2001) Beautiful faces have variable reward value: fMRI and behavioral evidence. Neuron 32(3):537– 551.

14. Bzdok D , et al. (2016) Left inferior parietal lobe engagement in social cognition and language. Neuroscience & Biobehavioral Reviews.

15. Rikandi E , et al. (2016) Precuneus functioning differentiates first-episode psychosis patients during the fantasy movie Alice in Wonderland. Psychological Medicine:1– 12.

16. Kawabata H & Zeki S (2004) Neural correlates of beauty. Journal of neurophysiology 91(4):1699– 1705.

17. Jacobs RH , Renken R , & Cornelissen FW (2012) Neural correlates of visual aesthetics–beauty as the coalescence of stimulus and internal state. PLoS One 7(2):e31248.

18. Jacobsen T , Schubotz RI , Höfel L , & Cramon DYv (2006) Brain correlates of aesthetic judgment of beauty. Neuroimage 29(1):276– 285.

19. Cattaneo Z , et al. (2013) The world can look better: enhancing beauty experience with brain stimulation. Social cognitive and affective neuroscience:nst165.

20. Lansdell H (1964) Sex differences in hemispheric asymmetries of the human brain.

21. McEwen BS & Milner TA (2017) Understanding the broad influence of sex hormones and sex differences in the brain. Journal of Neuroscience Research 95(1–2):24– 39.

22. Hu L & Iannetti GD (2016) Painful Issues in Pain Prediction. Trends in neurosciences 39(4):212– 220.

23. Price DD (2000) Psychological and neural mechanisms of the affective dimension of pain. Science 288(5472):1769– 1772.

24. Borgomaneri S , Gazzola V , & Avenanti A (2015) Transcranial magnetic stimulation reveals two functionally distinct stages of motor cortex involvement during perception of emotional body language. Brain Structure and function 220(5):2765– 2781.

25. de Borst A & de Gelder B (2016) Clear signals or mixed messages: inter-individual emotion congruency modulates brain activity underlying affective body perception. Social cognitive and affective neuroscience:nsw039.

26. De Gelder B , Snyder J , Greve D , Gerard G , & Hadjikhani N (2004) Fear fosters flight: a mechanism for fear contagion when perceiving emotion expressed by a whole body. Proceedings of the National Academy of Sciences of the United States of America 101(47):16701– 16706.

27. Engelen T , de Graaf TA , Sack AT , & de Gelder B (2015) A causal role for inferior parietal lobule in emotion body perception. cortex 73:195– 202.

28. Vartanian O & Skov M (2014) Neural correlates of viewing paintings: evidence from a quantitative meta-analysis of functional magnetic resonance imaging data. Brain and cognition 87:52– 56.

29. Blakemore S-J , Bristow D , Bird G , Frith C , & Ward J (2005) Somatosensory activations during the observation of touch and a case of vision–touch synaesthesia. Brain 128(7):1571– 1583.

30. Fiorio M & Haggard P (2005) Viewing the body prepares the brain for touch: effects of TMS over somatosensory cortex. European Journal of Neuroscience 22(3):773– 777.

31. Ellingsen D-M , et al. (2014) In touch with your emotions: oxytocin and touch change social impressions while others’ facial expressions can alter touch. Psychoneuroendocrinology 39:11– 20.

32. Gazzola V , et al. (2012) Primary somatosensory cortex discriminates affective significance in social touch. Proceedings of the National Academy of Sciences 109(25):E1657– E1666.

33. Rushworth MF , Buckley MJ , Behrens TE , Walton ME , & Bannerman DM (2007) Functional organization of the medial frontal cortex. Current opinion in neurobiology 17(2):220– 227.

34. Shenhav A , Botvinick MM , & Cohen JD (2013) The expected value of control: an integrative theory of anterior cingulate cortex function. Neuron 79(2):217– 240.

35. Nee DE , Kastner S , & Brown JW (2011) Functional heterogeneity of conflict, error, task-switching, and unexpectedness effects within medial prefrontal cortex. Neuroimage 54(1):528– 540.

36. Niendam TA , et al. (2012) Meta-analytic evidence for a superordinate cognitive control network subserving diverse executive functions. Cognitive, Affective, & Behavioral Neuroscience 12(2):241– 268.

37. Ridderinkhof KR , Nieuwenhuis S , & Braver TS (2007) Medial frontal cortex function: An introduction and overview. Cognitive, Affective, & Behavioral Neuroscience 7(4):261– 265.

38. Shackman AJ , et al. (2011) The integration of negative affect, pain and cognitive control in the cingulate cortex. Nature Reviews Neuroscience 12(3):154– 167.

39. Botvinick MM (2007) Conflict monitoring and decision making: reconciling two perspectives on anterior cingulate function. Cognitive, Affective, & Behavioral Neuroscience 7(4):356– 366.

40. Botvinick MM , Braver TS , Barch DM , Carter CS , & Cohen JD (2001) Conflict monitoring and cognitive control. Psychological review 108(3):624.

41. Botvinick MM , Cohen JD , & Carter CS (2004) Conflict monitoring and anterior cingulate cortex: an update. Trends in cognitive sciences 8(12):539– 546.

42. Critchley HD (2005) Neural mechanisms of autonomic, affective, and cognitive integration. Journal of Comparative Neurology 493(1):154– 166.

43. Rudebeck PH , et al. (2014) A role for primate subgenual cingulate cortex in sustaining autonomic arousal. Proceedings of the National Academy of Sciences 111(14):5391– 5396.

44. Skuse DH & Gallagher L (2009) Dopaminergic-neuropeptide interactions in the social brain. Trends in cognitive sciences 13(1):27– 35.

45. Zak PJ , Kurzban R , & Matzner WT (2004) The neurobiology of trust. Annals of the New York Academy of Sciences 1032(1):224– 227.

46. Freedman L & Cassell M (1994) Relationship of thalamic basal forebrain projection neurons to the peptidergic innervation of the midline thalamus. Journal of Comparative Neurology 348(3):321– 342.

47. Damasio AR , Everitt BJ , & Bishop D (1996) The somatic marker hypothesis and the possible functions of the prefrontal cortex [and discussion]. Philosophical Transactions of the Royal Society of London B: Biological Sciences 351(1346):1413– 1420.

48. Marusak H , et al. (2016) You say ‘prefrontal cortex’and I say ‘anterior cingulate’: meta-analysis of spatial overlap in amygdala-to-prefrontal connectivity and internalizing symptomology. Translational Psychiatry 6(11):e944.

49. Aleong R & Paus T (2010) Neural correlates of human body perception. Journal of Cognitive Neuroscience 22(3):482– 495.

50. Cazzato V , Mele S , & Urgesi C (2014) Gender differences in the neural underpinning of perceiving and appreciating the beauty of the body. Behavioural brain research 264:188– 196.

51. Kret ME , Pichon S , Grèzes J , & De Gelder B (2011) Men fear other men most: gender specific brain activations in perceiving threat from dynamic faces and bodies–an fMRI study. Frontiers in psychology 2:3.

52. Gallese V & Cuccio V (2014) The Paradigmatic Body: Embodied Simulation, Intersubjectivity, the Bodily Self, and Language. Open Mind, (Open MIND. Frankfurt am Main: MIND Group).

